# SARS-CoV-2 infects multiple species of North American deer mice and causes clinical disease in the California mouse

**DOI:** 10.1101/2022.08.22.504888

**Authors:** Juliette Lewis, Shijun Zhan, Allison C. Vilander, Anna C. Fagre, Hippokratis Kiaris, Tony Schountz

**Affiliations:** Department of Microbiology, Immunology and Pathology, Colorado State University, Fort Collins, CO; Drug Discovery & Biomedical Sciences, College of Pharmacy, University of South Carolina, Columbia, SC

## Abstract

Severe acute respiratory syndrome coronavirus-2 (SARS-CoV-2), the virus that causes coronavirus disease-19 (COVID-19), emerged in late 2019 in Wuhan, China and its rapid global spread has resulted in millions of deaths. An important public health consideration is the potential for SARS-CoV-2 to establish endemicity in a secondary animal reservoir outside of Asia or acquire adaptations that result in new variants with the ability to evade the immune response and reinfect the human population. Previous work has shown that North American deer mice (*Peromyscus maniculatus*) are susceptible and can transmit SARS-CoV-2 to naïve conspecifics, indicating its potential to serve as a wildlife reservoir for SARS-CoV-2 in North America. In this study, we report experimental SARS-CoV-2 susceptibility of two additional subspecies of the North American deer mouse and two additional deer mouse species, with infectious virus and viral RNA present in oral swabs and lung tissue of infected deer mice and neutralizing antibodies present at 15 days post-challenge. Moreover, some of one species, the California mouse (*P. californicus*) developed clinical disease, including one that required humane euthanasia. California mice often develop spontaneous liver disease, which may serve as a comorbidity for SARS-CoV-2 severity. The results of this study suggest broad susceptibility of rodents in the genus *Peromyscus* and further emphasize the potential of SARS-CoV-2 to infect a wide array of North American rodents.

**Importance:** A significant concern is the spillback of SARS-CoV-2 into North American wildlife species. We have determined that several species of peromyscine rodents, the most abundant mammals in North America, are susceptible to SARS-CoV-2 and that infection is likely long enough that the virus may be able to establish persistence in local rodent populations. Strikingly, some California mice developed clinical disease that suggests this species may be useful for the study of human co-morbidities often associated with severe and fatal COVID-19 disease.

## INTRODUCTION

Severe acute respiratory syndrome coronavirus-2 (SARS-CoV-2), the virus that causes Coronavirus disease-19 (COVID-19), emerged in late 2019 in Wuhan, China with rapid global spread resulting in millions of deaths (1). Molecular evidence suggests the zoonotic origin of SARS-CoV-2 is horseshoe bats of the genus *Rhinolophus,* which carry many SARS-related coronaviruses (2). COVID-19 remains an ongoing pandemic with hundreds of millions of cases and substantial global economic impacts.

The long-term pandemic status of COVID-19 is partially facilitated by the periodic emergence of new variants of SARS-CoV-2 that overcome immunity from vaccination or prior infection. The emergence of the omicron variant of SARS-CoV-2 in November 2021 sparked a new wave of infections globally due to its increased ability to infect convalescent and vaccinated individuals (3). The rapid accumulation of spike protein mutations that led to the omicron variant has led to the hypothesis that its progenitor may have acquired mutations during adaptation to a non-human host, perhaps rodents (4, 5). The potential for new variants to emerge because of adaptation to non-human hosts must be considered, and therefore identification and characterization of prospective non-human hosts is necessary to ensure preparedness.

Several mammalian species have been shown to be susceptible to infection with SARS-CoV-2, including tree shrews, Syrian hamsters, ferrets, cats, Egyptian fruit bats, mink, North American deer mice, and several species of non-human primates (6–13). Of note was the human-to-mink transmission of SARS-CoV-2 on mink farms in several countries, which resulted in localized human outbreaks of a mink-adapted SARS-CoV-2 variant (8). This exemplifies the need to examine animals living in close proximity to humans for their potential to act as hosts to SARS-CoV-2.

In our previous work, we determined that one subspecies of North American deer mice (*Peromyscus maniculatus nebrascensis*) is susceptible to experimental infection with SARS-CoV-2 and capable of transmission to naïve conspecifics for at least two passages. Another group reported similar findings in another subspecies of deer mouse (*P. m. rufinus*) (7). Deer mice are common peridomestic rodents that occupy a wide geographical range in North America (14, 15) and are the principal reservoir hosts of Sin Nombre orthohantavirus (16) and *Borrelia burgdorferi,* the causative agent of Lyme disease (17). Given their prominence in North American ecosystems, including on and around mink farms, it is feasible that SARS-CoV-2 could be introduced into wild deer mouse populations.

In this work, we expand the evidence of peromyscine rodent susceptibility to infection with a human isolate of SARS-CoV-2 to include two additional subspecies of the North American deer mouse group, including the Sonora white-footed mouse (*P. m. sonoriensis*) and the prairie deer mouse (*P. m. bairdii*). We also demonstrate susceptibility of two additional deer mouse species, the Oldfield mouse (*P. polionotus subgriseus*) and the California mouse (*P. californicus*), suggesting that rodents in genus *Peromyscus* are broadly susceptible to SARS-CoV-2. The examined species also expand the geographical range in which susceptible deer mice exist, increasing the risk of zooanthroponotic transmission of SARS-COV-2 from humans to wild rodents.

## MATERIALS AND METHODS

### Viruses

SARS-CoV-2 (isolate 2019-nCoV/USA-WA1, NR52281) was obtained from BEI Resources. Virus was passaged twice on Vero E6 cells (ATCC CRL-1586) in 2% FBS-DMEM containing penicillin and streptomycin at 37°C under 5% CO_2_ to generate stock virus used in these experiments. Virus was stored at −80° C. All work with infectious virus was performed at BSL-3 with approval from the Colorado State University Institutional Biosafety Committee.

### Animal procedures-multi-species study

All rodent work was approved by the CSU Institutional Animal Care and Use Committee (protocol 993). Deer mice were intranasally inoculated under inhalation of isoflurane anesthesia with 1×10^4^ (*P. bairdii*), 1.5×10^4^ (*P. sonoriensis*), or 2×10^4^ (*P. polionotus* and *P. californicus*) TCID_50_ SARS-CoV-2 (dose based on appropriate inoculation volume for body size). On days 3, 6 and 15 post-challenge, all animals were orally swabbed and weighed, and three inoculated mice of each species were euthanized by inhalation anesthesia and thoracotomy followed by cardiac exsanguination. Oral swabs and weights were taken for remaining mice on day 10 post-challenge. Control rodents were euthanized on day 0 using procedure described above. Necropsies were performed on all mice and samples (blood, lung) were collected for viral RNA detection (by RT-qPCR), virus isolation, and serology. Remaining whole carcasses were fixed in 10% neutral buffered formalin for histopathology and immunohistochemistry. For the California mouse study, all mice were intranasally inoculated under inhalation isoflurane anesthesia with 2×10^4^ TCID_50_ SARS-CoV-2 and four were euthanized as controls on either day 3 or 10 post-challenge. All mice were orally swabbed, weighed, and observed for clinical signs daily. Necropsies were performed on mice euthanized on days 3, 6, and 10 post-challenge, and tissues (blood, lung) were collected for viral RNA detection (by RT-qPCR), virus isolation, and serology. Remaining whole carcass was fixed in 10% neutral buffered formalin for histopathology and immunohistochemistry

### Sample collection and processing

Swabs were stored in 200 μL of viral transfer media (VTM) at −80° C until being processed and analyzed. Swabs were vortexed for 10 seconds followed by centrifugation at 1,500 rcf for 10 minutes to pellet any debris. Resulting supernatant was used in downstream assays. Tissues were stored in 500 μL of VTM and either flash frozen in LN_2_ and stored at −80° C until processing and analysis (multi-species experiment) or immediately processed following collection and stored at −80° C until analysis (California mouse experiment). Processing of tissue samples included homogenization on a TissueLyser II (Qiagen) at 50hz for 5 minutes followed by centrifugation at 10,000 rcf to pellet debris. The resulting supernatants were used in downstream analysis, including infectious virus isolation and RNA extraction for viral RNA quantification. Blood was collected in a serum separator tube, allowed to clot for 20-30 minutes, and then centrifuged at 20,000 rcf for 90 seconds. Serum was collected and stored at −80° C until analysis.

### TCID_50_ of tissues and swabs

Supernatant from swabs or lung homogenate was plated in triplicate on 96-well plates seeded with Vero E6 cells in a 10-fold dilution series from 10^0^ to 10^−3^. Samples that were above the limit of detection were re-plated in triplicate on 96-well plates seeded with Vero E6 cells in a 10-fold dilution series from 10^−2^ to 10^−5^. All plates were scored four days post-infection and titers were calculated using the Reed-Meunch method (18).

### Viral RNA in tissues and swabs

RNA was extracted from swab or tissue supernatant using the Mag-Bind Viral DNA/RNA 96 Kit (Omega Biotek) on the KingFisher FLEX System (ThermoFisher Scientific) according to manufacturer’s instructions. Reverse transcriptase quantitative PCR (RT-qPCR) was run using the GoTaq RT-qPCR System (Promega) with the IDT E Assay First Line Screening primers and probe (Cat # 10006804) on a LightCycler 96 System (Roche, Cat# 05815916001).

### Serum neutralization assay

Serum neutralization was performed starting at a 1:10 dilution with a 2-fold dilution series. An equal volume of SARS-CoV-2 containing 100 TCID_50_s was added (final serum dilution of 1:20) and incubated for 1 hr at 37° C. The mixture was plated on Vero E6 cells and scored for cytopathic effect after five days. The titer was the reciprocal of the greatest dilution that conferred 100% protection.

### ELISA

Recombinant SARS-CoV-2 nucleocapsid antigen in PBS was coated onto 96-well plates overnight at 4° C. Plates were washed 3x with PBS-0.5% TWEEN-20 and 3x with PBS between each step, then blocked with 0.5% gelatin in PBS for 30 min. Serum samples were diluted 2-fold in 0.5% BSA-PBS starting 1:100 and incubated for 2 hours at room temperature. Detection antibody, anti-*Peromyscus leucopus* IgG H&L-HRP (SeraCare), was diluted 1:1000 and incubated for 1 hour, followed by ABTS substrate. Plates were read at 405 nm and samples were considered positive if the OD was 0.200 above the mean of the negative control serum samples (uninfected deer mice).

### Histopathology and immunohistochemistry

After samples were collected for molecular analysis, the entire carcass was fixed in 10% neutral buffered formalin. Representative samples from the lungs, liver, heart, and the entire head (sagittal section) were collected and routinely processed for histopathology. Tissues were labeled with hematoxylin and eosin (H&E) for histologic analysis. Anti-SARS-CoV-2 immunohistochemistry was performed on embedded lung tissue from a representative subset of lung tissues at days 3 and 4 post-challenge, selected based on virus detection by RT PCR, using rabbit anit-SARS-CoV-2 nucleocapsid antibody [NP1189] (ProSci) as a primary antibody with hematoxylin counterstain.

## RESULTS

### Infectious virus was detected in each species of deer mouse

To investigate the susceptibility of different deer mouse species and subspecies, nine young, adult deer mice from each species or subspecies and of both sexes were intranasally challenged with SARS-CoV-2. Three unchallenged controls from each species group were euthanized and necropsied on day zero of the experiment. Three challenged mice from each species group were euthanized and necropsied on days 3, 6, and 15 post-challenge with one additional oral swab taken at day 10. Infectious virus was detected in 1/9 oral swabs from each species group on day 3 post-challenge, and viral RNA was detected in most lung samples from mice euthanized on days 3 and 6 post-challenge (Table 1, Supplementary Table 1). Viral RNA was detected in oral swabs of all challenged mice on day 3, and infectious virus was detected in oral swabs from 3/4 species on day 3 post-challenge (prairie deer mouse, California mouse, and Oldfield mouse). Three of the four species (prairie deer mouse, Sonora white-footed mouse, and Oldfield mouse) had detectable neutralizing antibodies on day 15 post-challenge, with an average geometric mean titer of 87 (SD ± 39). California mouse samples for this time point were compromised by mold contamination.

**Table 1:** Detection of viral RNA (vRNA) and infectious virus in lungs and oral swabs and serology. Values represented as number of positive samples out of total number of samples.

| Prairie deer mouse |  |  |  |  |  |
| --- | --- | --- | --- | --- | --- |
| days post-challenge | vRNA in lung | infectious virus in lung | vRNA in oral swab | infectious virus in oral swab | neutralizing antibodies |
| 3 | 3/3 | 2/3 | 9/9 | 1/9 | 0/3 |
| 6 | 3/3 | 1/3 | 6/6 | 0/6 | 0/3 |
| 10 | N/A | N/A | 3/3 | 0/3 | N/A |
| 15 | 0/3 | 0/3 | 0/3 | 0/3 | 2/3 |
| Sonora white-footed mouse |  |  |  |  |  |
| Days post-challenge | vRNA in lung | infectious virus in lung | vRNA in oral swab | infectious virus in oral swab | neutralizing antibodies |
| 3 | 2/3 | 0/3 | 9/9 | 1/9 | 0/3 |
| 6 | 2/3 | 1/3 | 5/6 | 0/6 | 0/3 |
| 10 | N/A | N/A | 1/3 | 0/3 | N/A |
| 15 | 1/3 | 0/3 | 0/3 | 0/3 | 2/3 |
| Oldfield mouse |  |  |  |  |  |
| Days post-challenge | vRNA in lung | infectious virus in lung | vRNA in oral swab | infectious virus in oral swab | neutralizing antibodies |
| 3 | 3/3 | 0/3 | 9/9 | 1/9 | 0/3 |
| 6 | 3/3 | 1/3 | 4/6 | 0/6 | 0/3 |
| 10 | N/A | N/A | 3/3 | 0/3 | N/A |
| 15 | 0/3 | 0/3 | 0/3 | 0/3 | 2/3 |
| California mouse |  |  |  |  |  |
| Days post-challenge | vRNA in lung | infectious virus in lung | vRNA in oral swab | infectious virus in oral swab | neutralizing antibodies |
| 3 | 3/3 | 0/3 | 9/9 | 1/9 | 0/3 |
| 6 | 3/3 | 1/3 | 4/5 | 0/6 | 0/3 |
| 10 | N/A | N/A | 1/2 | 0/3 | N/A |
| 15 | 0/2 | 0/2 | 0/2 | 0/3 | N/A |

### Clinical signs

No clinical signs were observed in the prairie deer mouse or Oldfield mouse at any timepoint in the study. The Sonora white-footed mice lost weight over the 15 day challenge study, despite a lack of other clinical signs (Figure 1). On day 3 post challenge, two California mice (1 female, 1 male) were observed to have ruffled fur, a hunched posture, and lethargic behavior. When the California mice were examined on day 4, the male appeared to have improved whereas the female was moribund and was humanely euthanized for necropsy. This mouse had hepatomegaly and prominent hepatic reticular pattern. It was suspected that the mouse was suffering from hepatic lipidosis (or, in humans, non-alcoholic fatty liver disease (NAFLD)). Three other California mice had a grossly prominent hepatic reticular pattern and hepatomegaly upon necropsy, including the male that displayed clinical signs on day 3. California mice lost significantly more weight 3 days post-challenge than any other group. To further investigate California mice as a model for SARS-CoV-2 disease, additional California mice were infected with SARS-CoV-2. Serial necropsies were performed on days 3, 6, and 10 post-challenge. No clinical signs were observed in these mice for the duration of the study.

**Figure 1.**
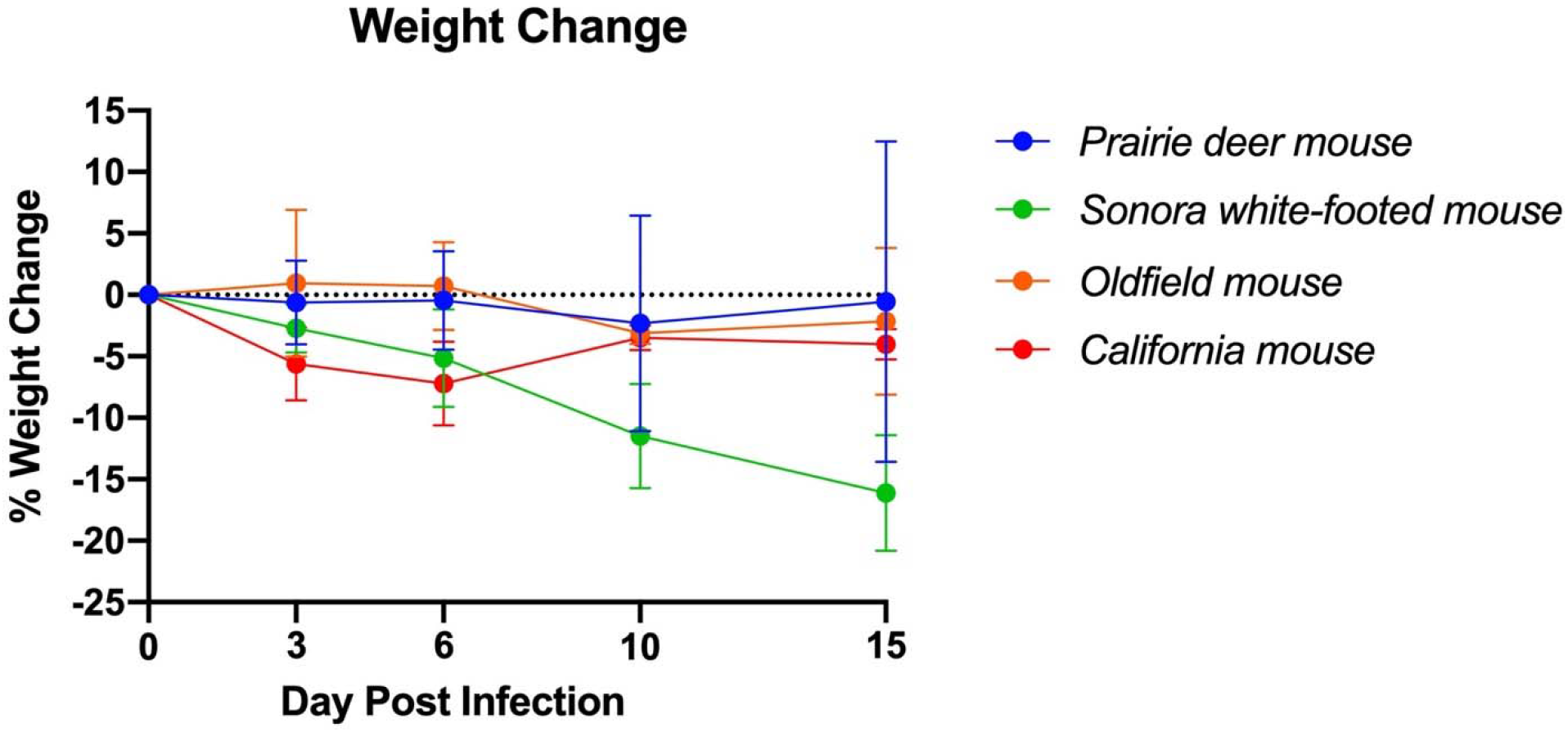
Weight change of deer mouse species during SARS-CoV-2 infection. On day 3 post-challenge, the California mice had significantly greater weight loss than the prairie deer mouse (*p*=0.004), Sonora white-footed mouse (*p*=0.027), and Oldfield mouse (*p*=0.010). At 6 days post-challenge, the California mouse had significantly greater weight loss than the prairie deer mouse (*p*=0.016) and the Oldfield mouse (*p*=0.005).

### California mice become infected with SARS-CoV-2

We further developed California mice as an animal model of SARS-CoV-2 infection following our observation of clinical disease in two mice during our initial investigation. Our follow-up study indicated that the mice became infected, with viral RNA detected through day 10 post-challenge in oral swabs and lungs (Figures 2A, 2C). Infectious virus was detected in oral swabs on days 1, 2, and 3 post-challenge (average 3.16 x 10^2^ ± 2.93 TCID_50_/mL, Figure 2B) and in all day 3 lungs (average 5.60 x 10^4^ ± 5.39 x 10^4^ TCID_50_/mL, Figure 2D). Serum IgG to nucleocapsid was detected in all mice at 10 days post-challenge (geometric mean titer 400, 95%CI [180,887] Figure 2E), and neutralizing antibody was detectable in 5 of 6 mice (geometric mean titer 92, 95% CI [45,189], Figure 2F). The SARS-CoV-2 challenged group showed significantly greater weight loss than the uninfected controls at days 1 and 3 post-challenge (Figure 3). At day 10 the control mice had a significant decrease in weight compared to infected controls. The cause of the weight loss is unknown. As the infected mice had all cleared detectible infectious virus by day 10, the weight loss in the control group at this time point was not significant to the conclusions of this study.

**Figure 2.**
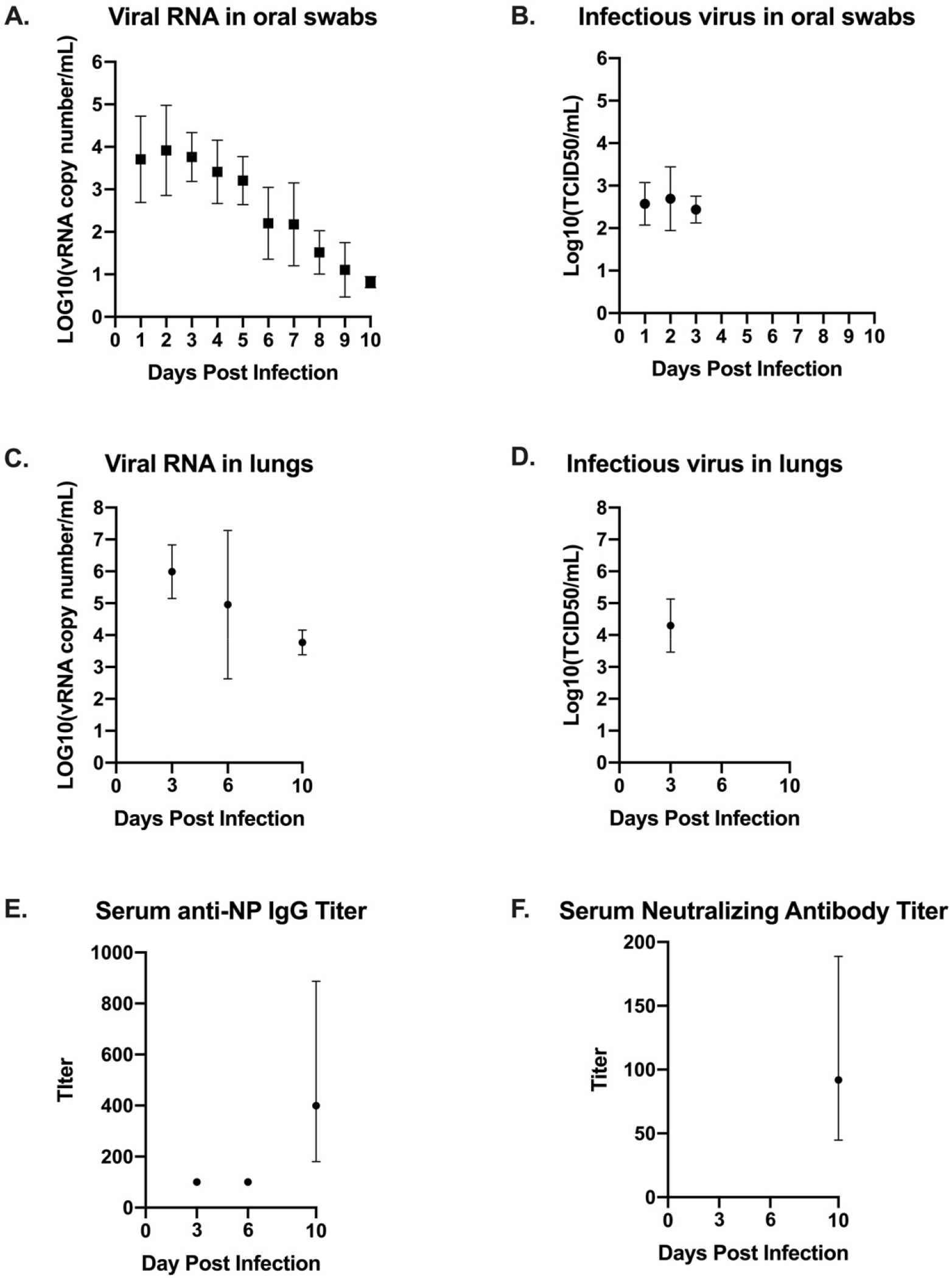
Virus detection and serology for California mice (*P. californicus insignis)*. Oral swabs collected daily (days 0-10 post-challenge) were tested for viral RNA (A) and infectious virus (B). Lung homogenate collected from mice euthanized on days 3, 6, and 10 post-challenge were also tested for viral RNA (C) and infectious virus (D). Serum collected from mice on days 3, 6, and 10 post-challenge was evaluated for quantification of serum IgG determined by indirect ELISA (E) and serum neutralizing antibody titer determined by serum neutralization assay (F).

**Figure 3.**
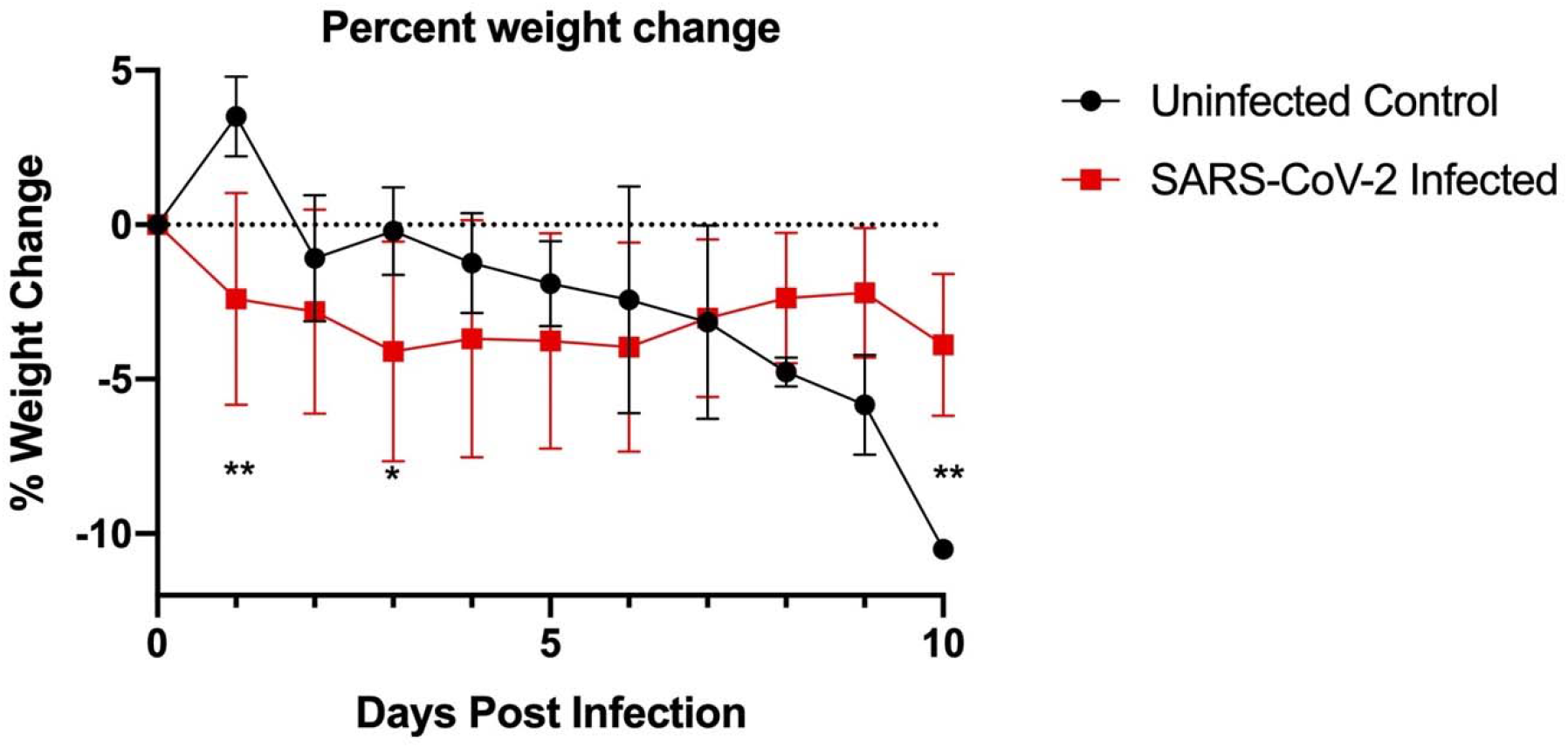
California mouse (*P. californicus insignis*) weight change during SARS-CoV-2 infection. SARS-CoV-2-challenged California mice had significantly greater weight loss on day 1 post-challenge (*p*=0.003) and day 3 post-challenge (*p*=0.047) as compared to the uninfected controls. The uninfected control group had significantly more weight loss than the challenged group at day 10 post-challenge (*p*=0.008).

### Histopathology and Immunohistochemistry

Lung, liver, spleen, nasal skin and mucosa, and heart from all mice were evaluated for pathology. The most severe inflammation was observed in the lungs and nasal passages in all species. The nasal skin and mucosa, heart, and liver did not show histologic differences in any of the groups. Lung was the most affected tissue and infected mice showed variable severity of interstitial lymphohistiocytic and fibrinous pneumonia with edema, reactive vascular endothelium with neutrophilic margination and perivasculitis, and rarely minimal to mild bronchial epithelial hyperplasia (Figure 4A– B). Necrosuppurative rhinitis was the second most common finding in infected mice (Figure 4C). Other observed pathology included mild lymphohistiocytic myocarditis, microvascular hepatic lipidosis, necrotizing and lymphohistiocytic dermatitis of the muzzle and lymphoplasmacytic perineuritis. Lesions in all species, when present, were most severe at day 3 post-challenge versus days 6/10/15. Pneumonia was present in all groups but was minimal to mild in the Sonora white-footed mouse.

Immunohistochemistry for anti-SARS-CoV-2 nucleocapsid had minimal rare positive labeling of bronchiolar epithelium in mice with virus detectible by RT-PCR (Figure 4D).

**Figure 4.**
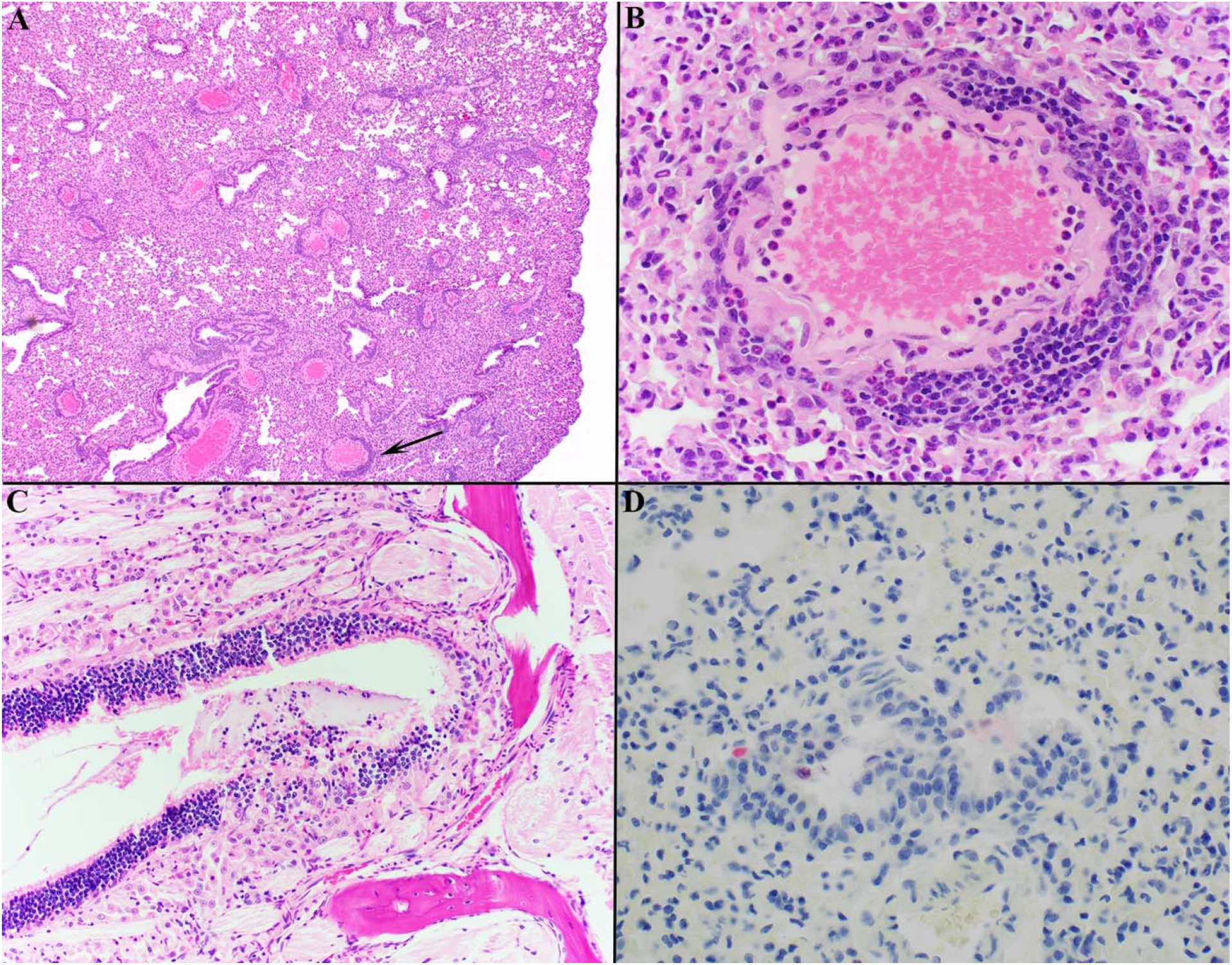
Histology of representative lesions in SARS-CoV-2 infected *Peromyscus spp*. Interstitial lymphohistiocytic pneumonia (A; Oldfield mouse) was the most common finding with multifocal vessels with segmental reactive endothelium, neutrophilic margination, and perivasculitis (A; Black arrow. Magnified in B). Necrosuppurative rhinitis was the second most common finding (C; California mouse). Anti-SARS-CoV-2 IHC shows rare positive labeling (red) of bronchial epithelial cells within the lungs (D; Sonora white-footed mouse). Hematoxylin counterstain. In six of the twelve California mice, there was hepatic lipidosis that ranged from mild to severe. The mouse that became moribund during the study demonstrated the most severe hepatic lipidosis histologically (Figure 5A). Lymphohistiocytic pneumonia in this mouse was minimal to mild but there was increased bronchial epithelial hyperplasia compared to other animals and this animal had the strongest anti-SARS-CoV-2 IHC labeling of the hyperplastic epithelium and scattered histiocytes (5B-C).

**Figure 5.**
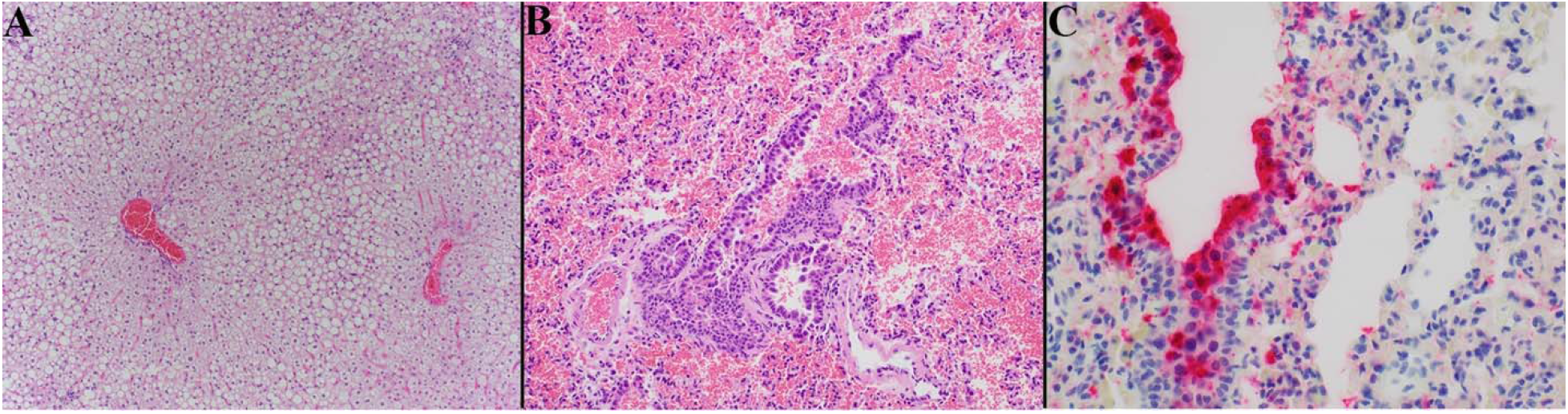
Photomicrographs of California mouse liver and pulmonary lesions. A single California mouse became moribund and at necropsy was found to have hepatomegaly. Histopathology showed severe diffuse distention of hepatocytes by lipid vacuoles consistent with hepatic lipidosis (A). The lungs of this animal exhibited only minimal to mild lymphohistiocytic pneumonia with moderate hyperplasia of the bronchiolar epithelium (B) but there was marked anti-SARS-CoV-2 positive labeling (red) of hyperplastic bronchial epithelium and histiocytes (C). Hematoxylin counterstain.

## Discussion

The North American genus *Peromyscus*, collectively known as deer mice, contains more than 50 species and has been studied extensively for their use as animal models of human disease and for their ability to host and transmit medically important pathogens (19). Many peromyscine rodents live in peri-domestic habitats, encountering humans through shared habitats as well as during outdoor activities (e.g. camping, field research). The experiments described here, along with previous work, confirm that three species of deer mice (the North American deer mouse, the Oldfield mouse, and the California mouse) and two subspecies of the North American deer mouse are susceptible to experimental SARS-CoV-2 infection with a human isolate of the virus (6, 7). There is some uncertainty as to the phylogenetic classification of the subspecies of deer mice used in this work, as it was proposed in 2019 by Greenbaum *et al.* that the North American deer mouse (*P. maniculatus*) be split into the eastern deer mouse (*P. maniculatus*) and western deer mouse (*P. sonoriensis*) (20). This taxonomic change would increase the number of confirmed susceptible deer mouse species to four.

Experimental infection resulted in asymptomatic infection in all species other than the California mouse, of which two individuals displayed clinical signs and one required euthanasia. Interestingly, the moribund California mouse that was euthanized early had significant immunostaining for SARS-CoV-2 nucleocapsid in the lungs but minimal histological signs of pneumonia, whereas asymptomatic animals in this study had more signs of pneumonia and very little immunostaining. This could indicate that immunosuppression or immunocompromise played a role in the infection outcome for that animal. For most animals, infection lasted through day 10 post-challenge, and by day 15 neutralizing antibodies were present in all species. The mostly asymptomatic and short-lived nature of SARS-CoV-2 infection in the observed species warrants investigation into their potential use as animal models for asymptomatic infection. The weight loss in some uninfected California mice was striking; however, this species is monogamous and we have observed loss of appetite after disruption of breeding pairs, particularly males, in the USC breeding colony. The separation of these animals after arrival at CSU may have contributed to weight loss. Additionally, the life span of deer mice, which can live up to 8 years in captivity vs ~2 years for laboratory mice (transgenic human ACE2) and Syrian hamsters (*Mesocricetus auratus*) (21) that are also used for SARS-CoV-2 studies, potentiates them as models for immune durability in vaccine and reinfection studies.

The California mouse has been studied as a model for several obesity-related conditions including type II diabetes mellitus and NAFLD (22, 23). California mice spontaneously develop these conditions when consuming a high-fat diet. In this study, we determined that California mice can be experimentally infected with SARS-CoV-2, with evidence of infectious virus in the lungs, oral shedding, and seroconversion. Both California mice that exhibited clinical signs in this study had hepatic lipidosis as determined by blinded pathological review, indicating that obesity-related conditions may have similar consequences in California mice as in humans during SARS-CoV-2 infection. NAFLD has been shown to predict COVID-19 progression and severity and COVID-19-related liver injury in human patients (24, 25). Therefore, the California mouse may be a suitable comorbidity model for NAFLD or other obesity related conditions and SARS-CoV-2. A larger study using California mice that have been fed a high-fat diet could address questions about liver dysfunction and SARS-CoV-2 disease, along with metabolic testing of the animals to confirm the conditions.

Peromyscine rodents and Syrian hamsters belong to the family Cricetidae. Deer mice and Syrian hamsters develop similar disease; however, California mice are the first small animal model to develop significant disease upon SARS-CoV-2 infection, other than transgenic mice that express human ACE2. This study used the WA1 isolate of SARS-CoV-2, thus future studies should examine other variants of concern, particularly delta variants that can cause more severe disease in humans. Considering their long lifespans, their outbred nature, availability of annotated genomes for several species, ready availability, and easy laboratory management, peromyscine rodents may serve as superior models to existing small animal models used in SARS-CoV-2 studies.

## Acknowledgements

This work was supported by grants from the National Institute of Allergy and Infectious Disease (R01 AI140442) and the National Science Foundation (2033260).

## Supplementary Materials

**Supplementary Table 1.** P-values of multiple unpaired T-tests performed between species groups. Group notation is as follows: *P. m. bairdii* (A), *P. m. sonoriensis* (B), *P. polionotus* (C), *P. californicus* (D). Comparisons with P<0.05 were considered significant.

| day post-challenge | A vs B | A vs C | A vs D | B vs C | B vs D | C vs D |
| --- | --- | --- | --- | --- | --- | --- |
| 3 | 0.128 | 0.509 | 0.004 | 0.101 | 0.027 | 0.010 |
| 6 | 0.068 | 0.610 | 0.016 | 0.023 | 0.393 | 0.005 |
| 10 | 0.179 | 0.884 | 0.871 | 0.029 | 0.089 | 0.694 |
| 15 | 0.124 | 0.857 | 0.744 | 0.034 | 0.043 | 0.703 |

